# Contribution of opsins and chromophores to cone pigment variation across populations of Lake Victoria cichlids

**DOI:** 10.1101/2021.07.09.451730

**Authors:** Elodie Wilwert, Rampal S. Etienne, Louis van de Zande, Martine E. Maan

## Abstract

Adaptation to heterogeneous sensory environments has been implicated as a key parameter in speciation. Cichlid fish are a textbook example of divergent visual adaptation, mediated by variation in the sequences and expression levels of cone opsin genes (encoding the protein component of visual pigments). In some vertebrates including fish, visual sensitivity is also tuned by the ratio of Vitamin A_1_/A_2_-derived chromophores (i.e. the light-sensitive component of the visual pigment, bound to the opsin protein), where higher proportions of A_2_ cause a more red-shifted wavelength absorbance. Here, we explore variation in chromophore ratios across multiple cichlid populations in Lake Victoria, using as a proxy the enzyme CYP27C1 that catalyses the conversion of Vitamin A_1_-into A_2_. We focus on sympatric *Pundamilia* cichlids, where species with blue or red male coloration co-occur at multiple islands, but occupy different depths and consequently different visual habitats. In the red species, we found higher *cyp27c1* expression in populations from turbid-water than from clear-water locations, but there was no such pattern in the blue species. Across populations, differences between the sympatric species in *cyp27c1* expression had a consistent relationship with species differences in opsin expression patterns, but the red/blue identity reversed between clear- and turbid-water locations. To assess the contribution of heritable versus environmental causes of variation, we tested whether light manipulations induce a change in *cyp27c1* expression in the laboratory. We found that *cyp27c1* expression was not influenced by experimental light conditions, suggesting that the observed variation in the wild is due to genetic differences. Establishing the biological importance of this variation requires testing the link between *cyp27c1* expression and A_1_/A_2_ ratios in the eye, as well as its consequences for visual performance.

## 1 Introduction

Local adaptation of sensory traits can initiate or strengthen species divergence. This is because sensory traits are important for both ecological performance and sexual communication (Boughman 2002; Endler and Basolo 1998). There are compelling reasons to study the visual system in this context: it is a crucial determinant of fitness in many taxa and highly diverse among species (Endler 1991; Fernald 1988; Marshall and Vorobyev 2003). Especially aquatic environments induce fine-scale visual adaptation in vision-dependent organisms. Owing to the variation in underwater light conditions, divergent selection on visual system properties can be strong (Kullander *et al*. 2014; Loew and McFarland 1990; Partridge *et al*. 1989) and visual adaptation to local light environments has been documented in numerous fish species (Bowmaker *et al*. 1994; Ehlman *et al*. 2015; Lythgoe *et al*.1994; Partridge *et al*. 1989; Shand *et al*. 2008). In particular, populations that occur in multiple distinct visual environments provide a good opportunity to explore variation in cone pigments within and between species. Here, we explore variation in cone pigments across populations of cichlid fish from different visual environments. Cichlids form one of the most species-rich families among vertebrates (Kocher 2004) and inhabit a broad range of visual environments (Shelly *et al*. 2006). They possess highly diverse visual systems (Carleton and Kocher 2001; Seehausen *et al*. 2008; Terai *et al*. 2017) and provide some of the best supported examples of speciation by divergent visual adaptation (Hofmann *et al*. 2009; Seehausen *et al*. 2008; Spady *et al*. 2005).

In fish, as in all vertebrates, visual information is captured by visual pigments in the eye, composed of an opsin protein covalently bound to a photosensitive Vitamin-A-derived chromophore. Visual sensitivity depends on the interaction between these components and may be tuned by variation in either component (Carleton *et al*. 2016). Cichlid fish possess several major opsin proteins: one rod opsin (RH1) for dim light vision and five cone opsins involved in colour vision (UV-sensitive (SWS1), blue-sensitive (SWS2), green-sensitive (Rh2a and Rh2b) and red-sensitive (LWS)). Variation in colour vision among cichlids comes from differences in the set of opsin genes that is expressed, from variation in opsin expression levels within that set, and from differences in opsin coding sequences (Carleton and Kocher 2001; Carleton *et al*. 2005; Carleton *et al*. 2008; Carleton *et al*. 2016; Hofmann *et al*. 2009; Larmuseau *et al*. 2009; Terai *et al*. 2006). This variation can be amplified by variation in chromophore composition (Saarinen *et al*. 2012; Sugawara *et al*. 2005; Terai *et al*. 2006; Torres-Dowdall *et al*. 2017). Two types of chromophores occur in fish visual pigments, derived from either Vitamin-A_1_- or Vitamin-A_2_. Higher proportions of Vitamin-A_2_ shift pigment absorption maxima to longer wavelengths, with a stronger effect in longer wavelength absorbing opsins (Govardovskii *et al*. 2000; Harosi 1994; Parry and Bowmaker 2000) (Figure 1a). Chromophore composition varies among species: marine fish and some freshwater fish possess solely A_1_-derived chromophores, while most freshwater fish carry only A_2_ or A_1_/A_2_ mixtures (Bridges and Yoshikami 1970; Morshedian *et al*. 2017; Provencio *et al*. 1992; Reuter *et al*. 1971; Toyama *et al*. 2008; van der Meer and Bowmaker 1995). In some species, chromophore ratios are phenotypically plastic, changing with environmental and/or life-history variables, such as migration, development, diet, season or temperature (Munz and McFarland 1977; Suzuki *et al*. 1984). For example, the migratory Coho salmon (*Oncorhynchus kisutch*) shows annual shifts in A_1_/A_2_ usage, changing from high proportions of A_1_ in the sea to high proportions of A_2_ when migrating into freshwater streams for spawning (Temple *et al*. 2006). In the non-migratory rudd, *Scardinius erythrophthalmus*, chromophore ratios covary with age, with older fish expressing higher A_2_ proportions (Bridges and Yoshikami 1970). Switches between the two types of chromophores can occur within a few weeks (Munz and McFarland 1977) and can, therefore, serve as a way to adjust to short-term changes in light conditions. In cichlids, only a few studies have explored variation in chromophore composition. They suggest that A_1_-based chromophores tend to dominate in species inhabiting clear waters (e.g. Lake Malawi cichlids) (Carleton *et al*. 2000; Parry *et al*. 2005; Sugawara *et al*. 2005; Torres-Dowdall *et al*. 2017), while species occupying turbid waters show higher usage of A_2_-based chromophores (Escobar-Camacho *et al*. 2019; Terai *et al*.. 2006), indicating that variation in A_2_-usage in cichlids may be important for perceiving long-wavelength light. Here, we explore the potential contribution of differential chromophore usage to visual adaptation in multiple cichlid populations from Lake Victoria.

**Figure 1.**
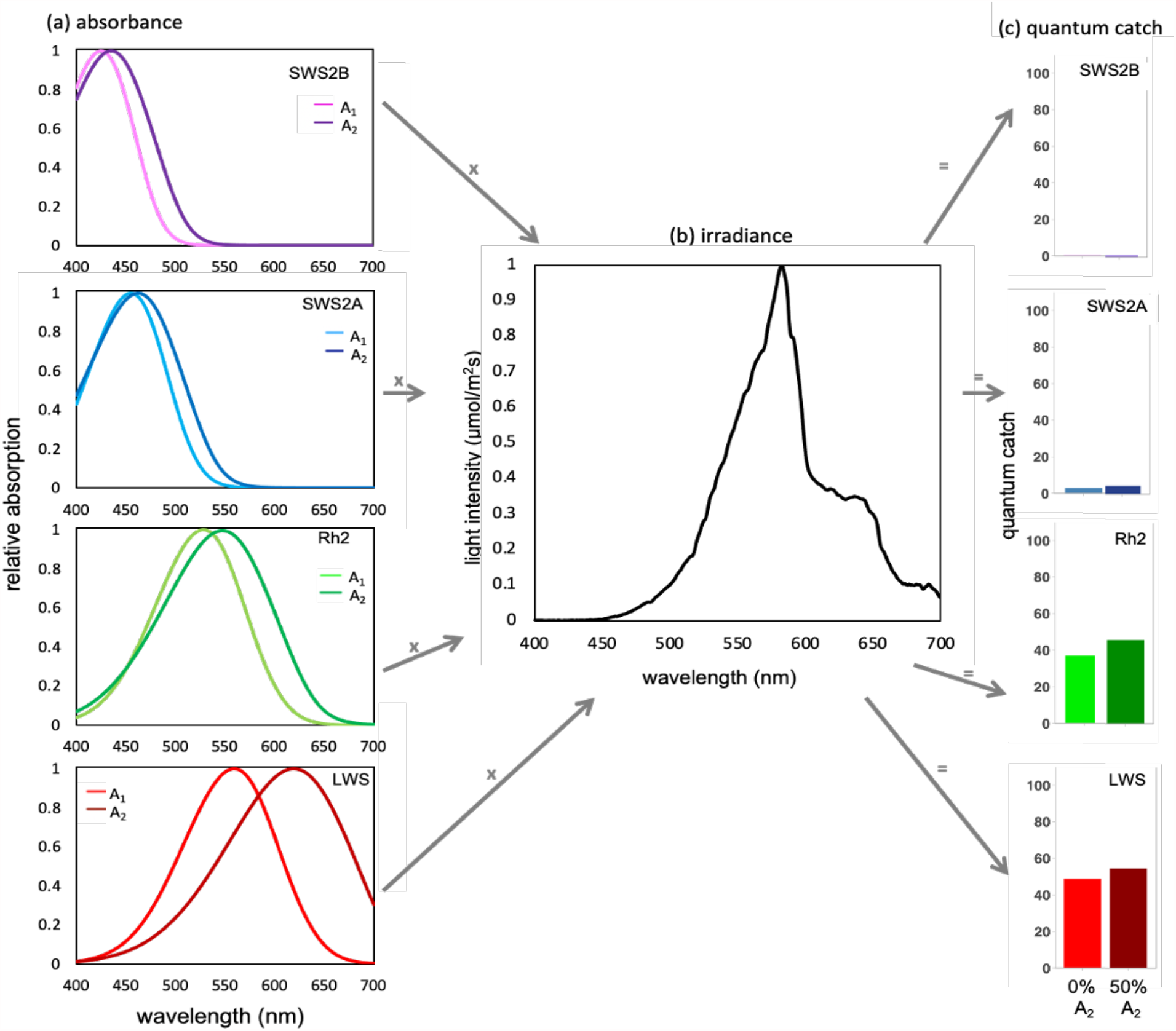
Schematic of the quantum catch calculation. a) Usage of A_2_ -derived chromophores instead of A_1_ - derived chromophores generates a shift in pigment peak sensitivity, with larger shifts for opsins with peak sensitivity at longer wavelengths. b) Pigment absorption curves, for either A_1_ or A_2_, are multiplied with the ambient light spectrum experienced by the focal population. As an example, depicted is the downwelling irradiance spectrum at Makobe island at 6m depth. c) The resulting quantum catch estimate for different A_1_ /A_2_ ratios for each opsin. Here illustrated for 0% and 50% A_2_.

We focus on closely related populations of *Pundamilia* cichlids (red and blue phenotypes) from several locations that differ in water clarity, and hence different light environments. Correlated differences in male coloration, female mate preference, photic environment and visual system properties suggest that visual adaptation to different light regimes contributes to *Pundamilia* divergence. However, chromophore usage has not been documented in *Pundamilia* and hence its role in divergent visual adaptation is unknown. Microspectrophotometry (MSP) of pigment absorption suggests that red phenotypes may use higher A_2_ proportions than blue phenotypes (Carleton *et al*. 2005).

Previous studies in a variety of vertebrate species have estimated chromophore ratios by MSP, or quantified alternative Vitamin A derivatives by high pressure liquid chromatography (HPLC). Enright *et al*. (2015) showed that in zebrafish, the conversion from Vitamin-A_1_ into A_2_ is mediated by the enzyme cytochrome p450 family 27 subfamily C polypeptide 1 (CYP27C1). In line with this, studies in bullfrog, zebrafish and lamprey have documented positive correlations between *cyp27c1* expression levels and A_2_ proportions in retinal pigments (Enright *et al*. 2015; Morshedian *et al*. 2017). This suggests that *cyp27c1* expression levels can be used as a proxy for A_2_ proportions. This is the approach we adopt in the present study. Specifically, we investigate in *Pundamilia* whether i) *cyp27c1* expression profiles vary between islands and phenotypes; ii) visual conditions with long-wavelength light are associated with higher expression levels of *cyp27c1*; iii) variation in *cyp27c1* expression is correlated with opsin expression patterns and iv) the observed patterns in *cyp27c1* and opsin expression optimize visual performance. Finally, given that chromophore usage may be influenced by both genetic and environmental factors, we v) explore the effect of different light regimes on *cyp27c1* expression levels in laboratory-reared fish.

## 2 Methods

### 2.1 Fish

#### Wild-caught fish

We included only male fish. We focus on *Pundamilia pundamilia* (Seehausen *et al*. 1998) and *Pundamilia nyererei* (Witte-Maas & Witte 1985) from the Speke Gulf and north-eastern Mwanza Gulf of Lake Victoria, and similar sympatric *Pundamilia* species pairs from the western and southern Mwanza Gulf of Lake Victoria (*P. sp*.*”pundamilia-like” and P. sp*.*”nyererei-like*”; Meier *et al*. 2017; Meier *et al*. 2018). Specifically, *Pundamilia* species were collected in 2014 at five rocky islands in south-eastern Lake Victoria; Mwanza Gulf: Luanso (−2.6889, 32.8842), Kissenda (−2.5494, 32.8276), Python (−2.6238, 35.8566) and Anchor (−2.5552, 32.8848); Speke Gulf: Makobe (−2.3654, 32.9228). Water transparency varies across islands, with more turbid waters at the southern end of the sampled region (i.e. Luanso, Kissenda and Python islands) and clearer waters at the northern end (i.e. Anchor and Makobe). Males of the species pair differ in nuptial coloration: *P. pundamilia* and *P. sp*.*”pundamilia-like”* display a blue/grey coloration and *P. nyererei* and *P. sp*.*”nyererei-like”* are yellow on the flanks and orange or red dorsally (Seehausen 1996). Females are inconspicuously coloured and exert colour-mediated assortative mate preferences (Haesler and Seehausen 2005; Seehausen and Van Alphen 1997; Selz *et al*. 2014; Stelkens *et al*. 2008). For simplicity, we write ‘blue phenotype’ for *P. pundamilia/P. sp*.*”pundamilia-like*” and ‘red phenotype’ for *P. nyererei/P. sp*.*”nyererei-like”*. Phenotypes tend to have different depth distributions coinciding with different photic environments: blue phenotypes inhabit shallow waters with broad-spectrum light, while red phenotypes occur at greater depths, where long-wavelength light (i.e. yellow and red) dominates (Seehausen *et al*. 2008). Until recently, all populations with blue males were classified as *P. pundamilia* and all populations with red males as *P. nyererei*. However, population genomic analyses revealed that populations from the southern and western Mwanza-Gulf (Kissenda and Python islands) represent a separate speciation event; they are therefore referred to as *P. sp*.*”pundamilia-like*” and *P. sp*.*”nyererei-like”* (Meier *et al*. 2017; Meier *et al*. 2018). At the most southern island, Luanso, blue and red phenotypes show no genetic differentiation (Meier *et al*. 2018) and fish were categorized into blue or red phenotypes by visually scoring coloration (as in Wright *et al*. 2019). Blue and red phenotypes have distinct visual system properties: they differ in the amino acid sequence of the long-wavelength sensitive opsin (LWS) and also show differences in opsin gene expression levels, corresponding to the differences in visual environment between geographic locations and depth ranges (Seehausen *et al*. 2008; Wright *et al*. 2019). In line with these differences, the red phenotype displays greater behavioural sensitivity to long wavelength light than the blue phenotype (at least in the most northern, clear-water location Makobe; Maan *et al*. 2006).

Sampling was conducted with permission of the Tanzanian Commission for Science and Technology (COSTECH-No. 2013-253-NA-2014-117). We collected 111 adult male fish by gillnetting and angling (Luanso = 10, Kissenda = 32, Python = 29, Anchor = 13, Makobe = 27). Capture depth was recorded for each individual. Fish were transported to the Tanzanian Fisheries Research Institute (TAFIRI - Mwanza Centre) and sacrificed by using 2-Phenoxyethanol (∼2.5ml/L) and subsequent cutting of the vertebral column. All fish were sacrificed in the early evening on the day of capture (17:00-20:00), to maximize RNA yield and minimize differences due to circadian variation in gene expression (Halstenberg *et al*. 2005; Yourick *et al*. 2019). Eyes were subsequently extracted, preserved in *RNAlater™* (Ambion) and frozen (−20°C).

#### Laboratory-reared fish

To explore the effects of light manipulation on *cyp27c1* expression levels, F1 and F2 offspring of wild-caught fish collected in 2010 at Python island were reared in light conditions mimicking the conditions experienced by each phenotype in their natural habitat at Python island (Figure S1). Fish were bred opportunistically with 18 dams and 15 sires. We used 75 male offspring resulting from thirty crosses (mother x father: 17 P x P; 21 N x N). *Pundamilia* are female mouthbrooders; eggs were removed from brooding females approximately 6 days after fertilization and divided evenly between the two light conditions. Fish were housed at 25±1°C on a 12L:12D light cycle and fed with commercial cichlid pellets and frozen Artemia, spirulina and krill. All specimens were sacrificed as adults, by applying an MS-222 (1 g/L) overdose and subsequent cutting of the vertebral column. Eyes were extracted, preserved in *RNAlater™* (Ambion) and frozen (−20°C). All fish were sacrificed in the late afternoon (16:00-18:00). This study was conducted under the approval of the Institutional Animal Care and Use Committee of the University of Groningen (DEC6205B; AVD105002016464).

### 2.2 Light measurements

In 2010, downwelling light intensity (µmol/m^2^* s) was measured at each island, using a BLK-C-100 spectrometer with an F-600-UV-VIS-SR optical fibre with CR2 cosine receptor (Stellar-Net, FL). Measurements were taken between 08:00 and 12:00 at depths down to 13 meters at 0.5 m increments. We took two independent series of measurements from Luanso, three from Kissenda and four from Makobe and Python island. Measurements were collected on different days, and we used the mean across sampling days for each depth measurement. Irradiance measurements were not conducted at Anchor island.

To explore the relationship between photic environments and fish visual system properties we calculated the orange ratio of each light spectrum. The orange ratio is the ratio of light transmitted in the 550-700nm range (yellow, orange and red) over the transmittance in the 400-549nm range (blue and green). As short wavelengths are more strongly absorbed and scattered with increasing depth in Lake Victoria, orange ratios increase with turbidity and depth (Figure S2). For each population, we calculated two measures of the orange ratio. First, population-level orange ratios were calculated for each population, based on depth distribution data from larger samples of fish (from Seehausen *et al*. 2008). Second, individual-level orange ratios were based on the capture depth of each individual fish that was sampled in the present study. Because no light measurements were available for Anchor island and prior work has shown that the water transparency at Anchor island is intermediate between Python and Makobe islands (Seehausen *et al*. 2008), we estimated the orange ratios at Anchor island as the medians of the ratios from Python and Makobe islands, following Wright *et al*. 2019.

### 2.3 Cyp27c1 gene expression

*Cpy27c1* expression was quantified using real-time quantitative polymerase chain reaction (qPCR). We removed the retina from the preserved eyes and extracted total RNA using Trizol (Ambion) followed by a DNase treatment to remove genomic DNA. RNA was reverse transcribed into cDNA using Oligo(dT)_18_ primer (Thermo Fisher Scientific) and RevertAid H Minus (Thermo Fisher Scientific) at 45°C. cDNA was diluted to a final concentration of 10ng/µl. Three housekeeping reference genes were used (HKGs): *ldh1, β-actin* and *gapdh2* (Jin *et al*. 2013; Torres -Dowdall *et al*. 2017). Stability of HKG expression was confirmed using RefFinder (Xie *et al*. 2012). qPCRs were run for 45 cycles (95°c for 3min, 95°C for 15 sec, 60°C for 25 sec, 72 for 30 sec) with specifically designed primers (Table S1) to amplify short fragments (200 bp). Each 20 µl reaction mixture contained 9µl gene-specific primer pairs, 1µl diluted cDNA sample and 10µl of SYBR Green PCR Master Mix (BioRad). Fluorescence was monitored on StepOnePlus Real-Time PCR System (Applied Biosystems ™StepOnePlus™Real-Time PCR System). To determine the critical threshold (Ct) and the initial concentration (N_0_) of *cyp27c1* and the three HKGs we used LinRegPCR (Ramakers and Ruijters 2003). Expression levels were based on two technical replicates. The following quality criteria were applied: PCR efficiency 1.75 – 2.25 and Ct standard deviation between duplicates ≤ 0.5. We used the following equation to calculate *cyp27c1* expression for each sample separately:

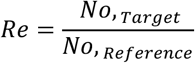

where N_0,Target_ is the initial concentration of *cyp27c1* and N_0,Reference_ is the geometric mean of the starting concentration of the three HKGs.

### 2.4 Opsin gene expression

To determine the relationship between opsin gene expression and *cyp27c1* expression in wild-caught fish, we used previously reported opsin gene expression data (i.e. SWS2b, SWS2a, Rh2 and LWS) from the same individuals from Wright *et al*. 2019.

### 2.5 Quantum catch estimates

To explore whether the observed variation between populations in *cyp27c1* expression (and opsin gene expression) enhances visual performance in the local light environment, we calculated quantum catch estimates (Qc), representing the number of photons captured by visual pigments in a given light environment. Quantum catches were estimated for the red and blue phenotypes at three locations (i.e. Kissenda, Python and Makobe islands). We excluded Luanso island because of low sample size and Anchor island because of the lack of light measurements. Quantum catches were calculated considering population-specific LWS genotype, with red phenotypes predominantly carrying LWS alleles conferring a more red-shifted sensitivity (H allele) than the blue phenotypes (P allele) (Seehausen *et al*. 2008), opsin expression profiles and depth ranges (i.e. visual environments), for three hypothetical A_1_ /A_2_ proportions (Figure 1). Quantum catches were calculated for each opsin using the following equation:

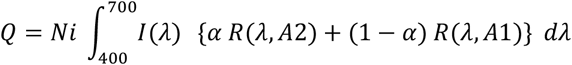

where *I*(*λ*) is the normalized irradiance spectrum at a specific capture depth and island, *N*_i_ is the relative opsin expression for each individual reported in (Wright *et al*. 2019), and R(*λ*) is the absorption spectrum of the visual pigments calculated for A_1_ and A_2_ separately (based on Govardovskii *et al*. 2000).

We used previously established peak sensitivities for each opsin in association with both A_1_ and A_2_-based chromophores (Carleton *et al*. 2005) (Table 1). To explore the impact of differential chromophore usage we estimated quantum catches for three hypothetical proportions of Vitamin A_2_ (designated by α): 10%, 30% and 50%. Quantum catch estimates were obtained using i) population-level irradiance spectra (based on the depth distributions of each population reported in (Seehausen *et al*. 2008) and ii) individual-level irradiance spectra (based on the individual capture depth of each fish).

**Table 1.** Peak sensitivities for each opsin and A_1_ and A_2_-based chromophores used for quantum catch calculations (Carleton et al. 2005).

| opsin | $\lambda_{\max}$ A1(nm) | $\lambda_{\max}$ A2 (nm) |
| --- | --- | --- |
| SWS2b | 455 | 462 |
| SWS2a | 425 | 435 |
| Rh2 | 528 | 547 |
| LWS (P-allele) | 544 | 604 |
| LWS (H-allele) | 559 | 619 |

### 2.6 Statistical analysis

All statistical analyses were performed in R (v 4.1.0; R Development Core Team 2021).

#### Wild-caught fish

*Cyp27c1* expression data were tested for outliers (1.5 x the interquartile range) separately for each population (i.e. phenotype-island combination). This resulted in *cyp27c1* expression data for 95 wild-caught fish (6 were excluded). After log transformation, we used linear models (LM) to explore if *cyp27c1* expression i) differed between islands and/or phenotypes, ii) covaried with water transparency, using the spectral midpoint from each island as an estimate for water transparency and iii) covaried with population-level and/or individual-level photic environment (i.e. orange ratio), as follows: *cyp27c1* expression ∼ island x phenotype + orange ratio. To determine the minimum adequate models we used stepwise backward selection using likelihood ratio tests (drop1 function, Crawley 2002). We used ANOVA to estimate parameters and *p*-values (car package, Fox *et al*. 2017). In case of more than two categories per fixed effect (i.e. island), we used post hoc Tukey tests (glht – multcomp package, Hothorn *et al*. 2008).

#### Laboratory-reared fish

*Cyp27c1* expression data were tested for outliers (1.5 x the interquartile range) separately for each phenotype and light treatment. This resulted in 66 laboratory samples for *cyp27c1* expression (9 removed). We tested for i) differences between wild-caught and laboratory-reared fish, using general linear models (GLM): *cyp27c1* expression ∼ phenotype x origin x light treatment, where ‘origin’ denotes wild-caught or laboratory-reared fish (i.e. including laboratory-reared fish from both light treatments) and light treatment denotes broad-spectrum light or red-shifted light; and ii) differences between light treatments, including only the laboratory-reared fish and using linear mixed effects modelling (lmer, package lme4) after log transforming the data: Relative gene expression ∼ phenotype x treatment + (1|mother ID) + (1|father ID), where treatment. To estimate parameter effects, *p*-values and degrees of freedom, we performed ’
sKRmodcomp’ (pbkrtest package; Halekoh and Højsgaard 2014)

## 3 Results

We found that *Pundamilia* from all five locations expressed *cyp27c1*, but expression levels differed between islands (χ^2^(4) = 3.23, *p* = 0.015). There was no overall difference between blue and red phenotypes (χ^2^(1) = 0.75, *p* = 0.387). Indeed, differences between islands were inconsistent between phenotypes, indicated by a significant island-by-phenotype interaction (χ^2^(4) = 4.12, *p* = 0.004) (Figure 2): *cyp27c1* expression decreased with water transparency in the red phenotypes (χ^2^(1) = 8.93, *p* = 0.003)), but not in the blue phenotypes (χ^2^(1) = 0.62, *p* = 0.429).

**Figure 2.**
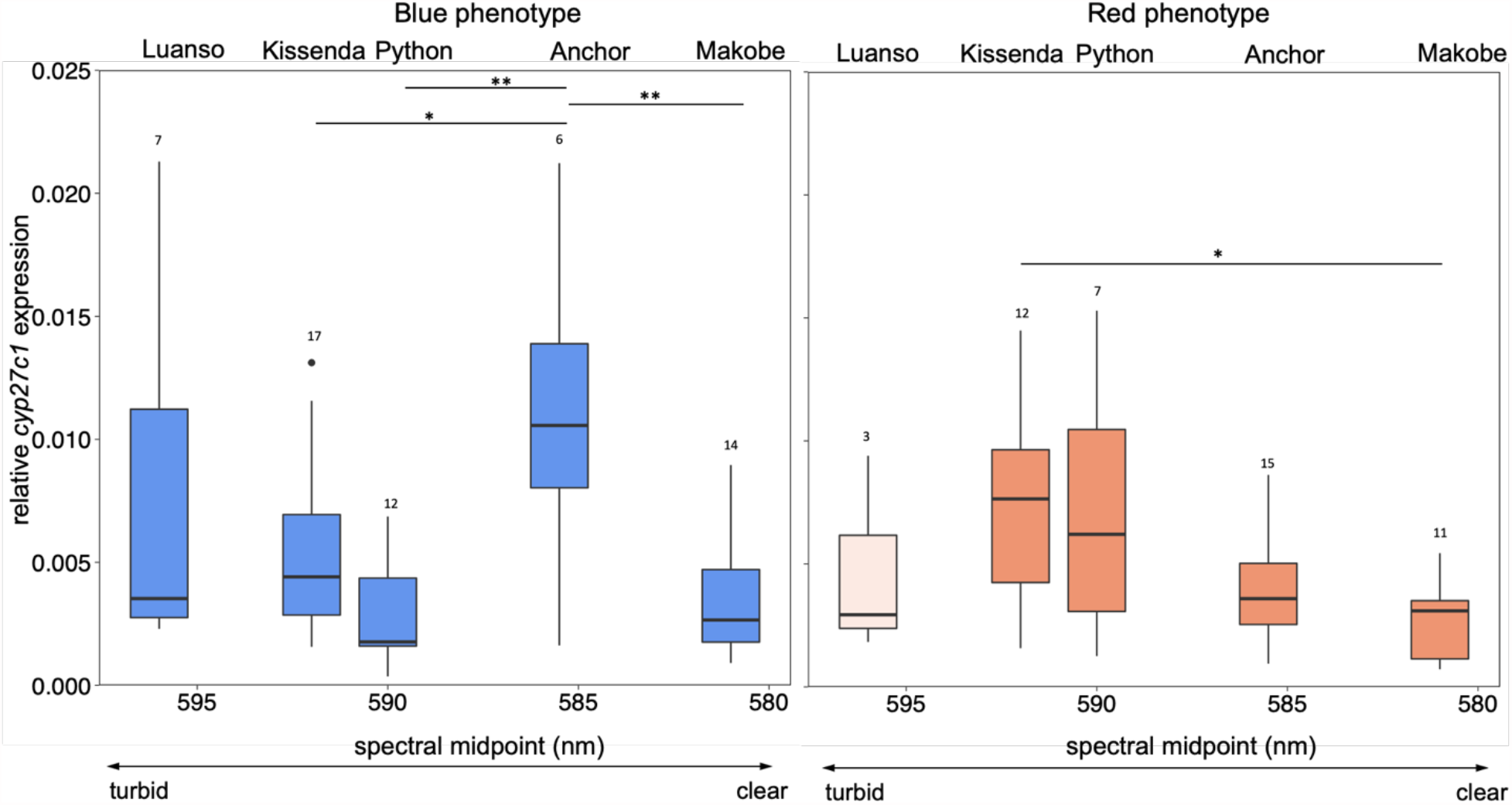
*Cyp27c1* expression in sympatric *Pundamilia* phenotypes at five locations. For both blue and red phenotypes, *cyp27c1* expression differed between islands. Geographic variation in expression levels was different between the red and blue phenotypes. Colours indicate phenotypes (Blue phenotype and Red phenotype). Red phenotypes from Luanso were rare (*n* = 3) and represented by a semi-transparent box. Boxes represent 25–75th percentiles, intercepted by the median. Error bars indicate 95% confidence intervals; black symbols are outliers. Sample sizes are given above each box. ** indicates *p* < 0.01, * indicates *p* < 0.05.

We then evaluated whether *cyp27c1* expression could be predicted by the local light environment. We found no evidence that *cyp27c1* expression covaried with population-level orange ratio (*χ*^*2*^(1) = 0.13, *p* = 0.716; Figure 3), nor with individual-level orange ratio (*χ*^*2*^(1) = 0.02, *p* = 0.898; Figure S3), indicating that the spectral composition of the local light environment did not affect *cyp27c1* expression levels.

**Figure 3.**
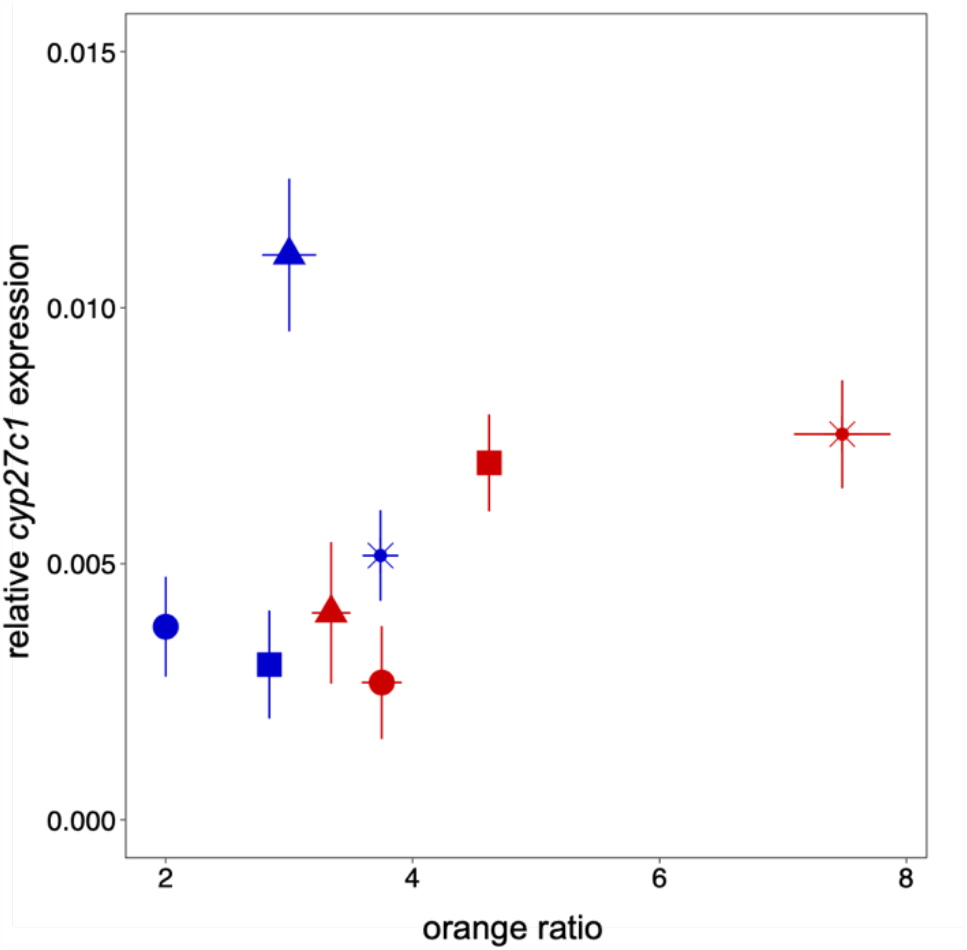
No covariation between *cyp27c1* expression and population-level orange ratio for *Pundamilia* populations at four locations. Symbols: Kissenda (**\***), Python (**▪**), Anchor (**▲**) and Makobe (**•**). Population-level orange ratios were derived from population-specific depth distributions presented in Seehausen *et al*. 2008. Colours indicate phenotypes (Blue phenotype and Red phenotype). Error bars represent ± standard error.

Regarding the within-island, between phenotype differences in *cyp27c1* expression, we found that sympatric blue and red phenotypes showed significant differences in *cyp27c1* expression at several islands, but the direction of the difference was inconsistent between islands (Figure 4a). At locations with higher turbidity (i.e. Kissenda and Python), *cyp27c1* expression tended to be higher in the red phenotypes, while at clear-water locations (i.e. Anchor and Makobe), *cyp27c1* expression tended to be higher in the blue phenotypes. Previous work in the same populations (Wright *et al*. 2019) reported that at clear-water islands (Makobe and Anchor) the red phenotypes expressed higher LWS (and lower Rh2) than the blue phenotypes, while at turbid-water locations (Python and Kissenda), the pattern was reversed (Figure 4b). In the present study, we observe that this reversal in Rh2/LWS expression is matched by a reversal in *cyp27c1* expression, suggesting that there is a consistent relationship between blue-red differences in *cyp27c1* expression and blue-red differences in opsin gene expression. In other words: at all locations, the phenotypes with the lower level of Rh2 expression (and higher level of LWS) tended to express lower levels of *cyp27c1*, but the identity of the phenotypes reversed between clear and turbid water locations (Figures 4 and 5). In contrast to this general consistency in the differences between sympatric blue and red phenotypes, we observed substantial individual variation (Figure 5), and no consistent relationships between *cyp27c1* and opsin gene expression at the individual level (Figure S4).

**Figure 4.**
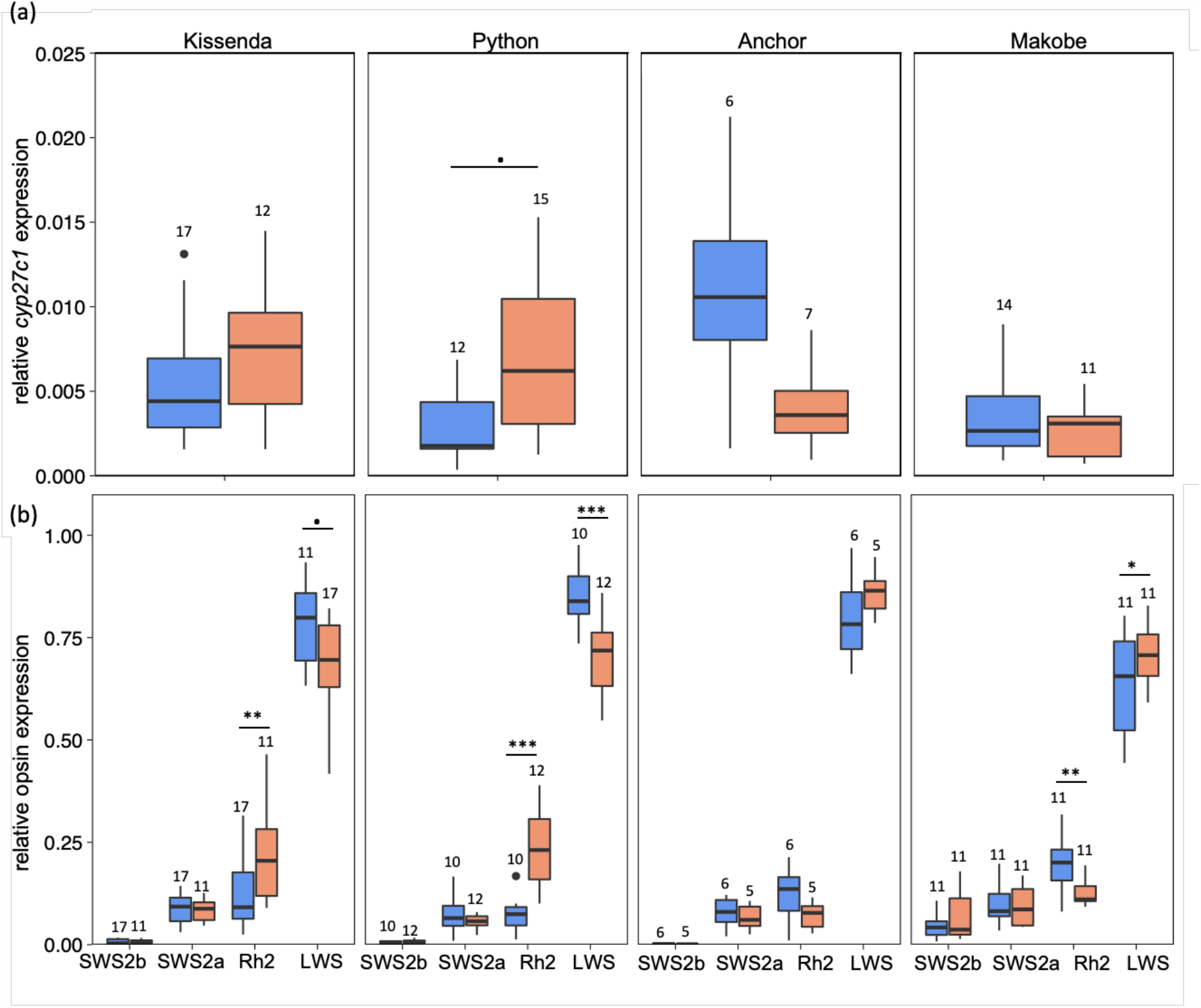
C*yp27c1* and opsin gene expression patterns in sympatric blue and red *Pundamilia* phenotypes from four locations. (a) *cyp27c1* expression levels. (b) opsin gene expression levels (from Wright *et al*. 2019). Boxes represent 25–75th percentiles, intercepted by the median. Error bars indicate 95% confidence intervals; black symbols are outliers. Sample sizes are indicated above each boxplot. ** indicates *p <* 0.001, ** indicates *p* < 0.01, * indicates *p* < 0.05 and • indicates *p* < 0.1.

**Figure 5.**
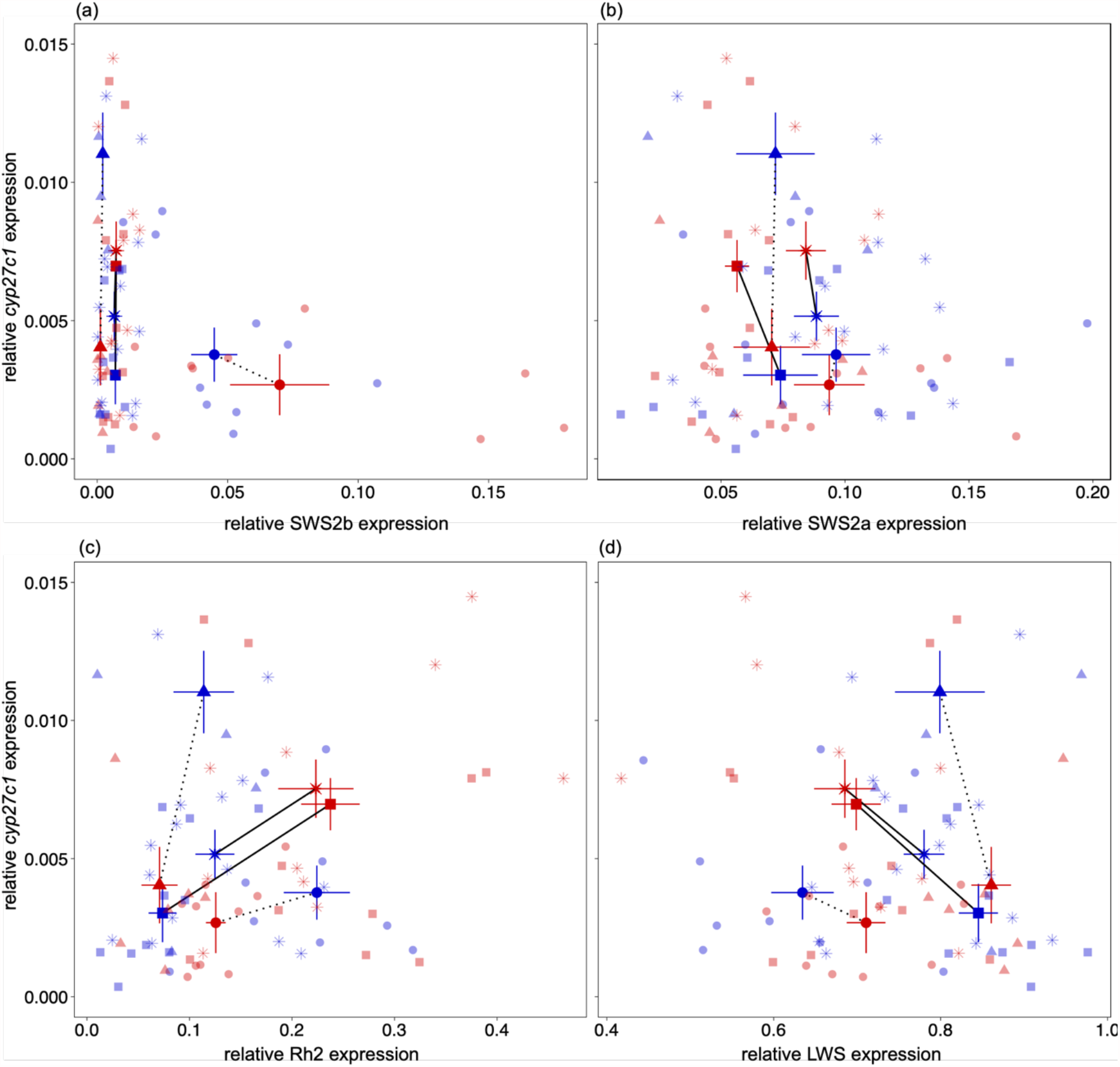
Relationships between *cyp27c1* and opsin gene expression in sympatric blue and red *Pundamilia* phenotypes from four locations. Colours indicate phenotypes (Blue phenotype and Red phenotype). Symbols indicate population means (Kissenda (**\***), Python (**▪**), Anchor (**▴**) and Makobe (**•**)). Shaded symbols indicate individual data points. Error bars represent ± standard error.

We used visual modelling to evaluate whether the observed patterns of *cyp27c1* expression maximize visual performance. Based on previous studies (Enright *et al*. 2015; Morshedian *et al*. 2017; Torres-Dowdall *et al*. 2017), we assume that higher levels of *cyp27c1* expression cause higher Vitamin A_2_ levels in the visual pigments and thereby red-shifted visual sensitivity. Thus, we predict that populations with higher *cyp27c1* expression levels, and hence higher A_2_ levels, obtain higher quantum catches (Qc) in red-shifted light conditions. The analysis did not support this hypothesis: a hypothetical increase in A_2_ proportions generates higher quantum catch estimates for every phenotype at each location (Figure 6). Quantum catch estimates based on individual depth ranges yielded similar results (Figure S5). Hence, these results do not provide evidence that the observed differences between populations in *cyp27c1* expression contribute to locally adapted visual performance.

**Figure 6.**
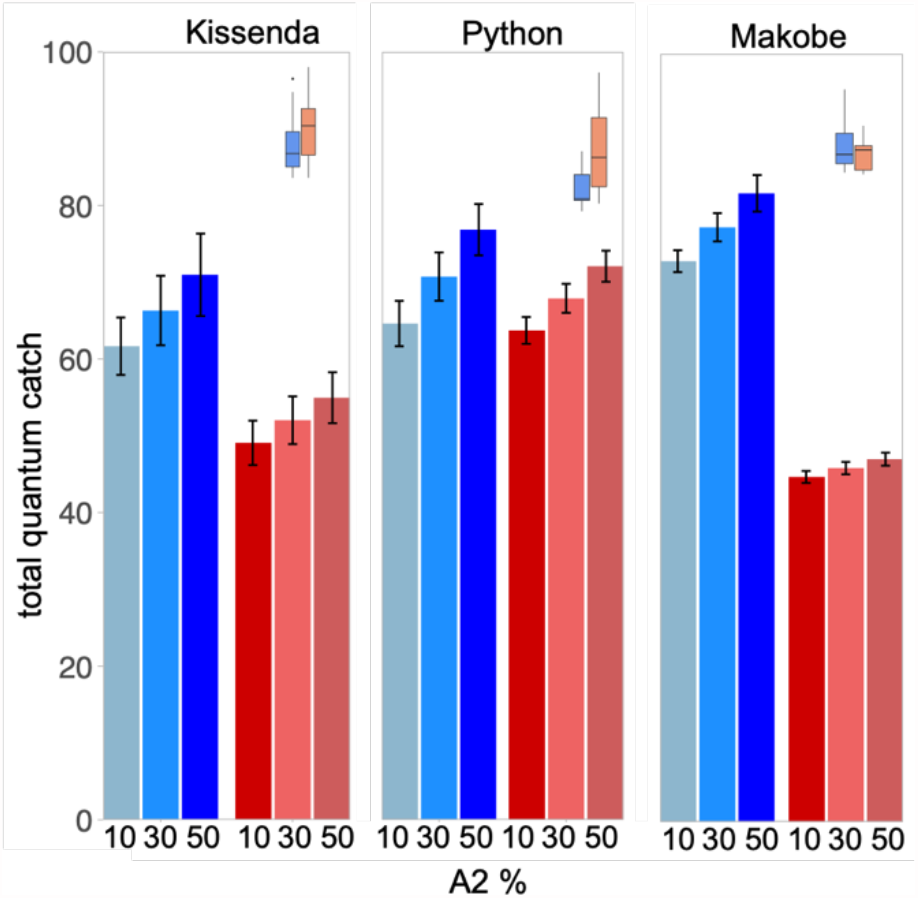
Variation in A_*2*_ proportion does not maximize visual performance for *Pundamilia* phenotypes at three locations. Bars represent different quantum catches for three hypothetical proportions of Vitamin A_2_ (i.e. 10%, 30% and 50%). Colours indicate phenotypes (Blue phenotype and Red phenotype) and error bars represent ± standard error. The top right panels show reported phenotype differences in *cyp27c1* expression (from Figure 4a).

To evaluate whether the observed variation in *cyp27c1* expression in the wild is due to genetic differences and/or phenotypic plasticity, we reared both *Pundamilia* phenotypes from one population (i.e. Python island) under two different light conditions, mimicking the light spectra in the natural shallow-water and deep-water environments of Python Island (i.e. broad spectrum light vs. red-shifted light spectrum). We found that overall, laboratory-bred individuals showed similar *cyp27c1* expression levels as their wild-caught counterparts (Figure 7a). However, there was a statistical trend indicating that the red-blue phenotype difference in the field (see above) was not maintained in the laboratory (phenotype by origin interaction: χ^2^(1) = 3.63, *p* = 0.05). Consequently, analysis including only laboratory-reared fish showed no difference between red and blue phenotypes (χ^2^(1) = 0.180, *p* = 0.675; Figure 7a). This was not explained by the different light conditions employed in the laboratory: we found no influence of the two rearing light conditions on *cyp27c1* expression (Figure 7b; *P. sp*.*“pundamilia*-like”: *z* = 1.33, *p* = 0.56; *P. sp*.*”nyererei*-like”: *z* = 0.03, *p* = 1.00). The difference between field and lab data was mostly due to the fact that laboratory-bred individuals of the red phenotype tended to have lower *cyp27c1* expression levels than their wild-caught counterparts, irrespective of the light conditions (*t* = -2.25; *p* = 0.05). In the blue phenotype, levels did not differ between laboratory-bred and wild-caught individuals (*t* = 2.18; *p* = 0.97).

**Figure 7.**
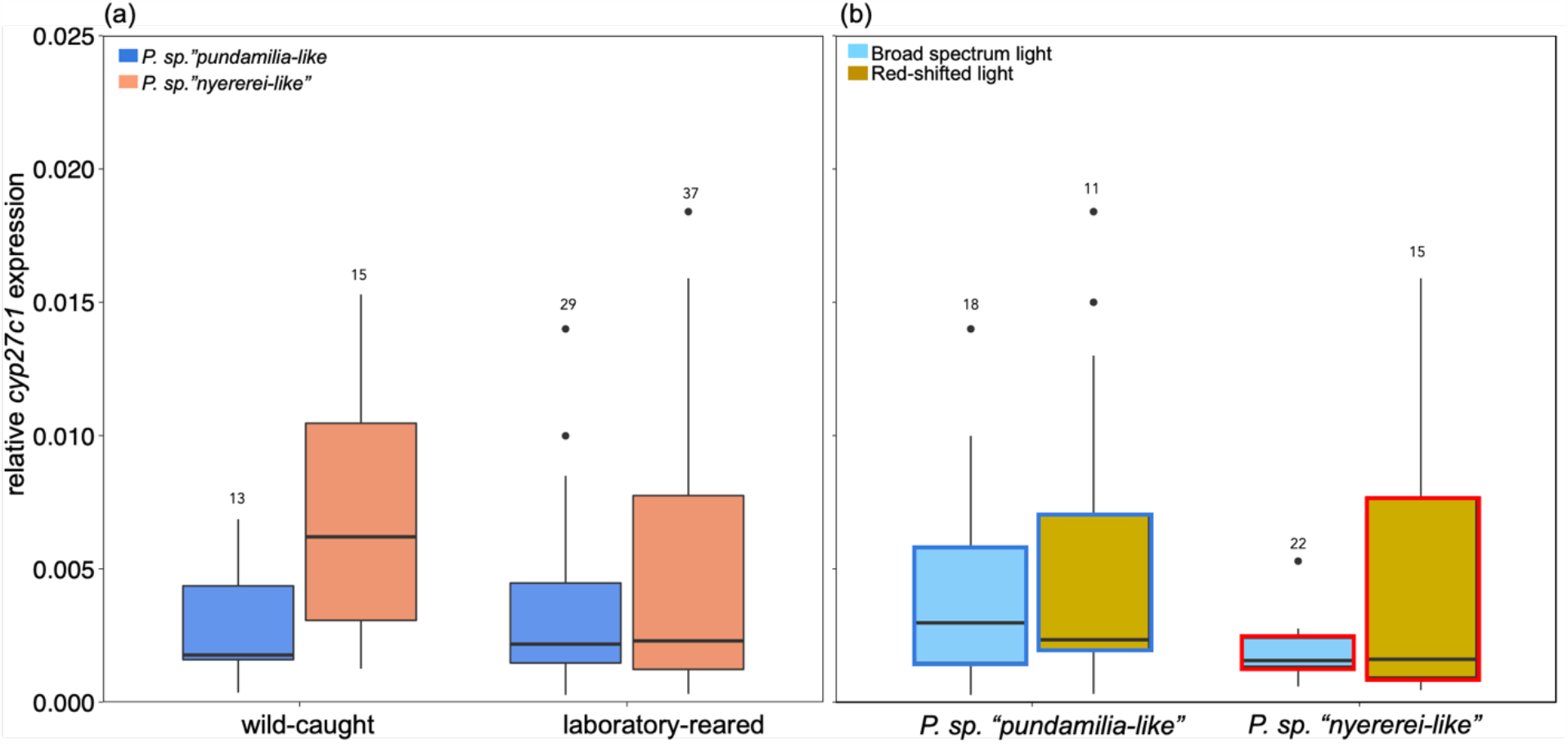
*Cyp27c1* expression in wild-caught and laboratory-reared fish. a) Cyp27c1 levels in laboratory fish were similar as in wild-caught fish b) Laboratory-reared phenotypes were not influenced by our rearing light conditions (broad-spectrum light versus red-shifted light). Boxes represent 25–75th percentiles, intercepted by the median. Error bars indicate 95% confidence intervals; black symbols are outliers. Sample sizes are indicated above each bar. •indicates *p* < 0.1.

## 4 Discussion

Cichlids are a major model system for visually mediated ecological speciation, based on evidence for divergent adaptation in opsin genes and opsin gene expression levels. In many vertebrates, particularly in fish, variation in chromophore usage also contributes to visual adaptation. This study provides the first investigation of differential chromophore usage across multiple cichlid populations from Lake Victoria, using as a proxy *cyp27c1*, an enzyme responsible to convert A_1_ into A_2_ –derived chromophores (Enright *et al*. 2015). We found that *Pundamilia* cichlids from five different islands express *cyp27c1* in the eye and that expression levels differ between populations. However, we also found substantial individual variation and no clear association between the visual environment and *cyp27c1* expression levels.

Typically, fish that inhabit red-shifted visual conditions (as in many turbid waters) use higher Vitamin A_2_ proportions (Bowmaker 1995). Yet, evidence for this pattern in cichlid species is mixed. Based on microspectrophotometry (MSP), it has been shown that cichlids from clear waters of Lake Malawi use only A_1_ (Carleton *et al*. 2000), while cichlids from turbid waters of Lake Victoria incorporate A_2_ (Carleton *et al*. 2005; Terai *et al*. 2006; van der Meer and Bowmaker 1995). However, indirect measurements of chromophore usage through *cyp27c1* expression are inconsistent. For example, Härer *et al*. (2018) observed that several closely related neotropical cichlid species expressed higher levels of *cyp27c1* in turbid waters (Lake Managua and Lake Nicaragua) than in clear waters (Lake Xiloa). However, in the Amazonian cichlid *Cichla monoculus* from Lake Gatun, fish captured in turbid waters expressed lower *cyp27c1* than fish from clear waters (Escobar-Camacho *et al*. 2019). In the present study, we observed that both *Pundamilia* phenotypes (i.e. red and blue) express *cyp27c1*, but we found no clear relationship with visual conditions (i.e. differences in water transparency across locations and variation in orange ratio across depths). Specifically, we found that the red *Pundamilia* phenotype displayed higher *cyp27c1* expression in turbid waters than in clear waters, but the blue phenotype showed no such pattern. A possible explanation for this species difference is the difference in depth distribution between the red and the blue phenotypes: the consistently shallow depth range of the blue phenotype entails a relatively consistent light environment across locations, while the red phenotypes occur in deeper waters where variation in water transparency has a much greater impact, generating larger variation in light conditions (Figure 3; Figure S7).

In various aquatic taxa including cichlids, species and population differences in opsin expression have been shown to correlate with variation in visual conditions (Carleton 2009; Carleton *et al*. 2016; Hofmann *et al*. 2009). Recent work in *Pundamilia* (Wright *et al*. 2019) demonstrated that at locations with higher turbidity (i.e. Kissenda and Python), the red phenotypes express more Rh2 and less LWS than the blue phenotypes, while this difference reverses at clear water locations (i.e. Anchor and Makobe). Visual modelling suggested that these patterns do not maximize quantum catch at each location, implying that the observed patterns could not be explained by local adaptation. Here, we explored whether *cyp27c1* expression could act as a compensatory mechanism and thereby explain these findings. Strikingly, we observed a similar reversal for *cyp27c1* expression levels across locations: at locations with higher turbidity (i.e. Kissenda and Python), *cyp27c1* expression tended to be higher in the red phenotypes than in the blue phenotypes, while at clear-water locations (i.e. Anchor and Makobe), we found the opposite. As a result, there is a consistent relationship between phenotype differences in *cyp27c1* expression levels and phenotype differences in opsin expression levels, where the phenotype expressing more Rh2 consistently also expresses more *cyp27c1* than its sympatric relative, but the red/blue identity reverses between clear-water and turbid-water locations. We propose three possible explanations for these findings. First, the observation that phenotypes with low LWS levels expressed high *cyp27c1* levels could reflect a compensatory mechanism, where reduced long-wavelength sensitivity (i.e. lower LWS expression) might be compensated with higher Vitamin-A_2_ usage (i.e. higher *cyp27c1* expression) and vice versa. However, our quantum catch estimates suggest that the observed variation does not maximize local visual performance, as any hypothetical increase in A_2_ proportion generates higher quantum catch estimates for every phenotype at each location. It should be noted that quantum catch estimates are a fairly crude measure of visual performance, taking into account only the ability to capture ambient light and ignoring more complex perceptual abilities such as colour discrimination and object recognition. Either way, a second explanation could be non-adaptive: the parallel reversal in opsin expression and *cyp27c1* expression between blue and red phenotypes may simply be a by-product of the evolutionary history of the study populations. Meier *et al*. (2017) found that the speciation event resulting in *P. pundamilia* and *P. nyererei* occurred outside the Mwanza Gulf, after which the species pair settled at Makobe island. Many generations later, *P. pundamilia* colonized the Western Mwanza Gulf (including Python island). After ∼1000 generations, *P. nyererei* from outside the Mwanza Gulf immigrated into the Western Mwanza Gulf and hybridized with the local *P. pundamilia* population, which resulted in a new speciation event in which *P. sp*.*”pundamilia-like”* and *P. sp*.*”nyererei-like”* emerged. Given this recent history, the characteristic expression patterns observed at Python and Kissenda, but not at Makobe and Anchor, may be a non-adaptive by-product of this event, rather than an adaptation to the visual environment at these locations. Third, in the last decades, intensive agriculture, deforestation and urban runoff have significantly increased nutrient loading causing the eutrophication of Lake Victoria (Scheren *et al*. 2000). This resulted in an increase in algal biomass, which in turn, together with silt carried by the rivers, decreased water transparency (Nyamewya *et al*. 2020). Hence, *Pundamilia* species have recently been exposed to environmental changes, allowing very little time to adapt and possibly explainning the lack of a relationship between opsin expression, *cyp27c1* expression and current visual conditions.

Rapid adjustments in visual system properties may improve individual visual performance in changing light conditions. Such plasticity may enhance population persistence and thereby provide a starting point for evolutionary change (Fusco and Minelli 2010; Price *et al*. 2003,West-Eberhard 2003). Several cichlid species have been shown to change opsin gene expression levels over development and in response to different light regimes (Härer *et al*. 2019; Nandamuri *et al*. 2018; Wright *et al*. 2020), suggesting a potential role of phenotypic plasticity in optimizing visual performance. Also in *Pundamilia*, expression levels of long-wavelength-sensitive (LWS) and short-wavelength-sensitive (SWS2a) opsins can be influenced by light manipulations (Wright *et al*. 2020). The present study is the first to explore plasticity in chromophore usage in *Pundamilia*. We observed that the phenotype difference in *cyp27c1* expression observed in the wild was reduced in laboratory-reared fish, which indicates a potential but modest contribution of plasticity to the variation observed in the wild. However, when reared under different light conditions in the lab, we did not detect an effect on *cyp27c1* expression levels. Together, these findings indicate that *cyp27c1* expression in *Pundamilia* is less plastic than opsin gene expression and that the observed variation in *cyp27c1* expression among natural populations largely reflects genetic differences. However, compared to other cichlid species, *cyp27c1* expression in *Pundamilia* seems to be expressed at very low levels (Torres-Dowdall *et al*. 2017), questioning the biological relevance of the observed variation in *cyp27c1* expression between phenotypes and locations.

A key assumption in this study was that *cyp27c1* expression and A_2_ proportion are positively correlated. However, only a few datapoints in neotropical cichlids are available to substantiate this assumption, with inconsistent results (Escobar-Camacho *et al*. 2019; Torres-Dowdall *et al*. 2017). Consequently, Escobar-Camacho *et al*. (2019) hypothesized that in cichlid fish, conversions between A_1_ and A_2_ might be regulated by other enzymes. This implies that further studies are needed to explore whether *cyp27c1* expression does actually reflect A_1_/A_2_ ratios in cichlid fish, and how this gene functions and interacts with other genes. We also need data from additional cichlid species, from different visual habitats, to evaluate the contribution of chromophore usage to cichlid visual adaptation.

## Acknowledgments

The authors thank the Tanzanian Commission for Science and Technology for research permission and the Tanzanian Fisheries Research Institute for hospitality and facilities. We thank Daniel Shane Wright for collecting the wild-caught samples and performing RNA isolation. We thank Roel van Eijk and Jolien Gay for helping to design the qPCR protocol. We thank Sjoerd Veenstra, Brendan Verbeek, Lucia Irazábal-González and Willem Diderich for taking care of the fish in the laboratory.

## Supporting information

**Figure S1**. Light conditions used in the laboratory

**Figure S2**. Orange ratios at study locations

**Table S1**. Gene specific primers for *cyp27c1* and reference genes

**Figure S3**. Relationship between *cyp27c1* expression and orange ratio based on the capture depths of the individuals used in this study

**Figure S4**. The relationships between *cyp27c1* expression and relative opsin gene expression

**Figure S5**. Quantum catch calculations based on individual capture depth

**Figure S7**. Variation in underwater light environments experienced by blue and red phenotypes at different sampling locations

## Author Contributions

MEM designed the study, together with RSE, LVDZ and EW. EW designed the qPCR protocol for *cyp27c1* and completed the laboratory work. EW performed the analysis, with assistance from MEM, RSE and LVDZ. EW wrote the manuscript with contributions of MEM, RSE and LVDZ. All authors approved the contents of this manuscript.

