## Supplementary information for "Contribution of opsins and chromophores to cone pigment variation across populations of Lake Victoria cichlids"

**Supporting Information**

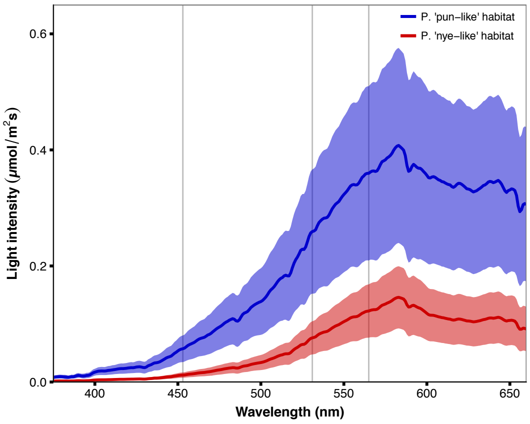


**Figure S1. Light conditions used in the laboratory.** Broad-spectrum and red-shifted rearing light conditions mimicking the natural environments of blue (i.e. broad spectrum light) and red (i.e. red-shifted light spectrum) phenotypes at Python island (from Maan *et al*. 2017). Vertical lines indicate the peak sensitivities of *Pundamilia* photoreceptors: SWS2a (453nm), Rh2 (531nm) and LWS(565nm) (Carelton *et al.* 2005).

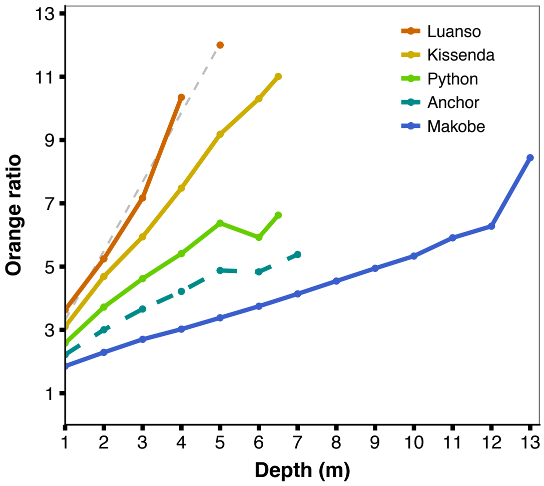


**Figure S2. Orange ratios at study locations.** The most turbid location, Luanso island, has the most red-shifted light spectrum and hence the highest orange ratios. Irradiance spectra for Anchor island were unavailable, so orange ratios were estimated as the median of the orange ratios of Python and Makobe islands. At Luanso island irradiance measurements were only available down to 4m depth, thus linear regression (dotted grey line) was used to estimate the orange ratio at 5m depth. Figure from Wright *et al.* (2019).

**Table S1. Gene specific primers for *cyp27c1* and reference genes.** Specificity of amplification was checked by Sanger sequencing (GATC Biotech) of purified PCR products.

| Gene | Forward Primer (5´-3´) | Reverse Primer (5´-3´) | Reference |
|---|---|---|---|
| *Cyp27c1* | GATGACTTGGTAGTTGGTGGA | GAACTTCTGAATGAGCCTTATGAG | This paper |
| *Gapdh2* | TGTCTTCCAGTGTATGAAGCC | GGTCGTATTTGTCCTCATTAACTC | This paper |
| *ldh2* | TTGGAGGTTTTAAGGAAAAGG | CAGGAACAAGGTGACGGTGGT | Torres-Dowdall *et al.* 2017 |
| *ß-actin* | AATCGTGCGTGACATCAAGG | AGTATTTACGCTCAGGTGGG | Jin *et al.* 2013 |

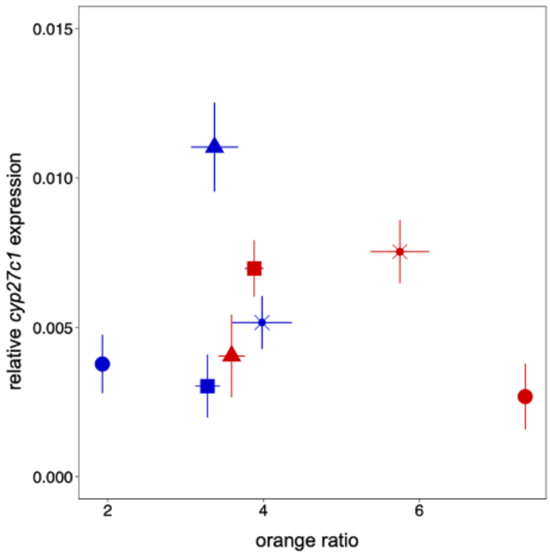


**Figure S3. Relationship between *cyp27c1* expression and orange ratio based on the capture depths of the individuals used in this study.** *Cyp27c1*expression does not increase with individual-level orange ratio across islands. Colours indicate phenotypes (Blue phenotype and Red phenotype) and shapes indicate islands (Kissenda ( * ), Python (■), Anchor (▲) and Makobe (●) ). Error bars represent ± standard error.

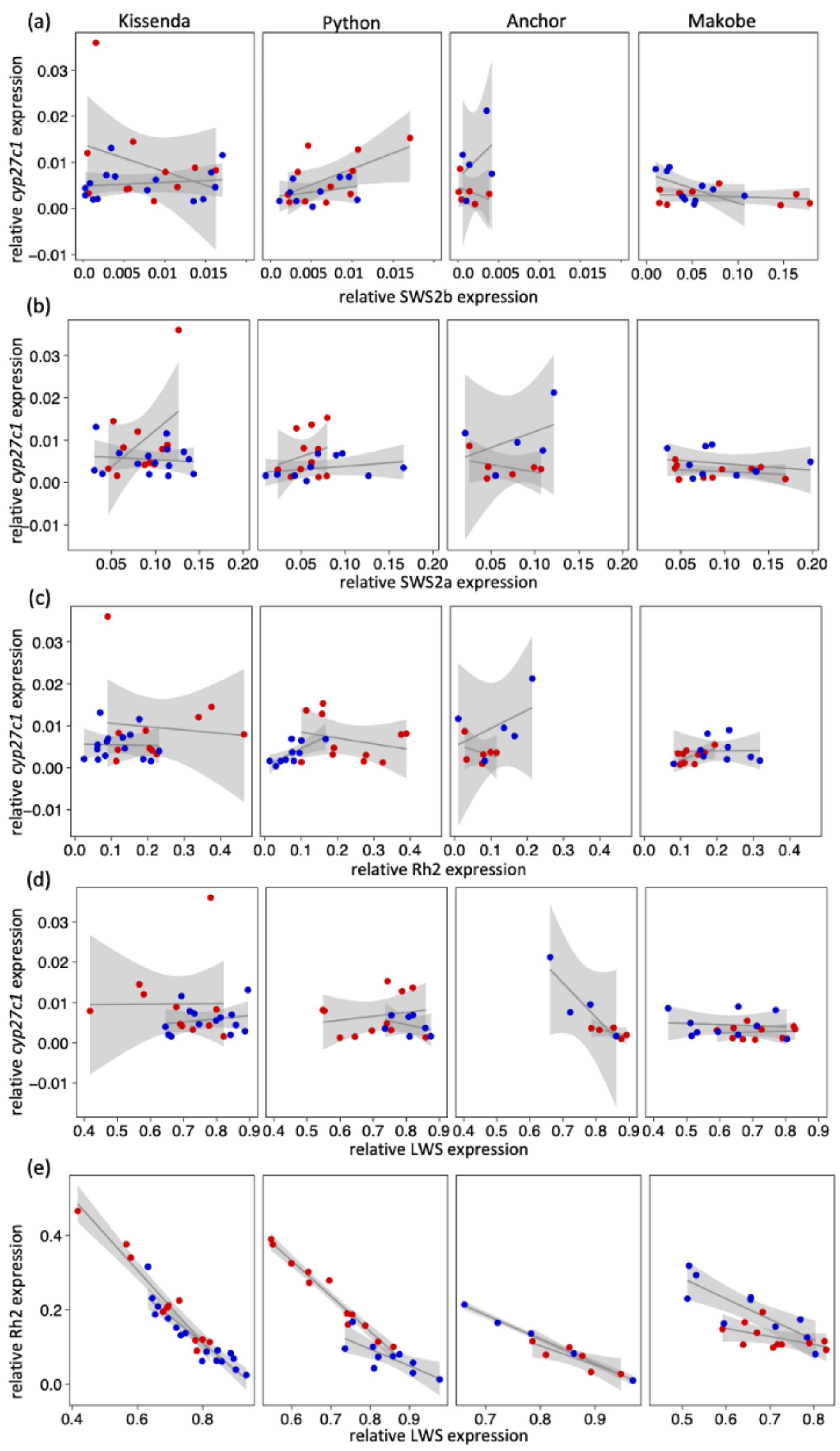


**Figure S4.** The relationships between *cyp27c1* expression and relative expression of a) SWS2b, b) SWS2a, c) Rh2 and d) LWS varied with location and phenotype. e) LWS and Rh2 expression are negatively correlated in all populations. Each symbol represents one individual, colours indicate phenotypes (Blue phenotype and Red phenotype) and shaded areas represent ± standard error.

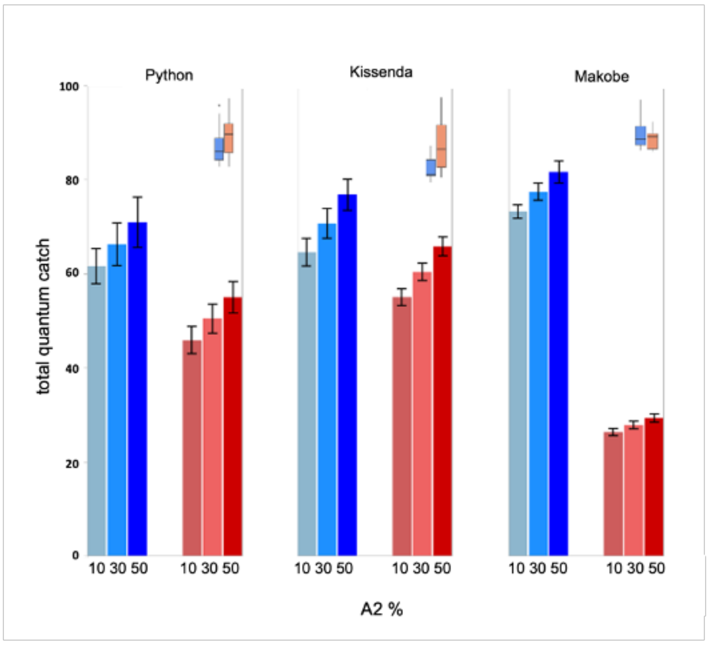


**Figure S5. Quantum catch calculations based on individual capture depth.** A hypothetical increase in $A_2$ proportion results in an increase in quantum catch estimates for each population ($p<0.001$). Bars represent different quantum catches for three different hypothetical proportions of Vitamin $A_2$ (i.e. 10%, 30% and 50%). Colours indicate phenotype (Blue phenotype and Red phenotype) and error bars represent ± standard error. The top right panels show reported phenotype differences in *cyp27c1* expression (see Figure 3a).

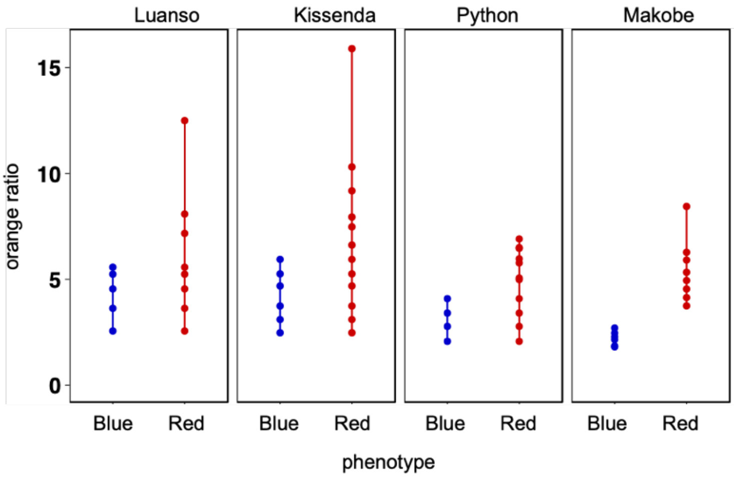


**Figure S7. Variation in underwater light environments experienced by blue and red phenotypes at different sampling locations.** Each symbol represents the orange ratio at a specific water depth (in 0.5m increments) where the phenotypes occur. Colours indicate phenotype (Blue and Red phenotype). Note that the maximum orange ratios for the red species represent the maximum gillnetting depths used in this study (the depth range of the red species extends to even greater depths, with higher orange ratios) while the maximum orange ratios for the blue species correspond to their actual depth distributions.
